# Ocular conjunctival inoculation of SARS-CoV-2 can cause mild COVID-19 in Rhesus macaques

**DOI:** 10.1101/2020.03.13.990036

**Authors:** Wei Deng, Linlin Bao, Hong Gao, Zhiguang Xiang, Yajin Qu, Zhiqi Song, Shunran Gong, Jiayi Liu, Jiangning Liu, Pin Yu, Feifei Qi, Yanfeng Xu, Fengli Li, Chong Xiao, Qi Lv, Jing Xue, Qiang Wei, Mingya Liu, Guanpeng Wang, Shunyi Wang, Haisheng Yu, Xing Liu, Wenjie Zhao, Yunlin Han, Chuan Qin

**Affiliations:** Key Laboratory of Human Disease Comparative Medicine, Chinese Ministry of Health, Beijing Key Laboratory for Animal Models of Emerging and Remerging Infectious Diseases, Institute of Laboratory Animal Science, Chinese Academy of Medical Sciences and Comparative Medicine Center, Peking Union Medical College, Beijing, China; Department of Radiology, Bejing Anzhen Hospital, Capital Medical University, Beijing, China

## Abstract

The outbreak of Corona Virus Disease 2019 caused by the severe acute respiratory syndrome coronavirus (SARS-CoV-2) is highly transmitted. The potential extra-respiratory transmission routes remain uncertain. Five rhesus macaques were inoculated with 1×10^6^ TCID_50_ of SARS-CoV-2 via conjunctival (CJ), intratracheal (IT), and intragastric (IG) routes, respectively. Remarkably, the CJ inoculated-macaques developed mild interstitial pneumonia and viral load was detectable in the conjunctival swabs at 1 days post-inoculation (dpi). Only via IT inoculation, viral load was detected in the anal swab at 1-7 dpi and macaque showed weight loss. However, viral load was undetectable after IG inoculation. Comparatively, viral load was higher in the nasolacrimal system but lesions of lung were relatively mild and local via CJ inoculation compared with that via IT inoculation, demonstrating distinct characteristics of virus dispersion. Both the two routes affected the alimentary tract. Therefore the clinicians need to protect eye while working with patients.

## Introduction

Corona Virus Disease 2019 (COVID-19) is highly infectious and transmitted mainly through human-to-human transmission via respiratory droplets when direct or close contact with the patients with SARS-CoV-2. The other potential transmission routes remain to be further researched. In some clinical cases, samples of tears and conjunctival secretions from both SARS-CoV^1^ and SARS-CoV-2 patients with conjunctivitis^1^ displayed detectable viral RNA. A previous study reported the case of a clinician who was infected with SARS-CoV-2 while working with patients under all safeguards except eye protection ^3^. By contrast, no SARS-CoV-2 could be detected by RT-PCR in 114 conjunctival swabs samples from patients with COVID-19 pneumonia^4^. The potential extra-respiratory portals that SARS-CoV-2 enter the host need to be further research by laboratory-confirmation for providing significant data to oversight and prevention for healthcare workers.

## Results

Five rhesus macaques between the ages of 3 and 5 years were inoculated with 1×10^6^ TCID_50_ of SARS-CoV-2 via three routes by ocular conjunctival inoculation (CJ-1 and CJ-2), intragastrical inoculation (IG-1 and IG-2), and intratracheal inoculation (IT-1) regarded as a comparison to compare the distributions and pathogenesis of viruses after enter the host via different routes (Figure 1).

**Fig. 1.**
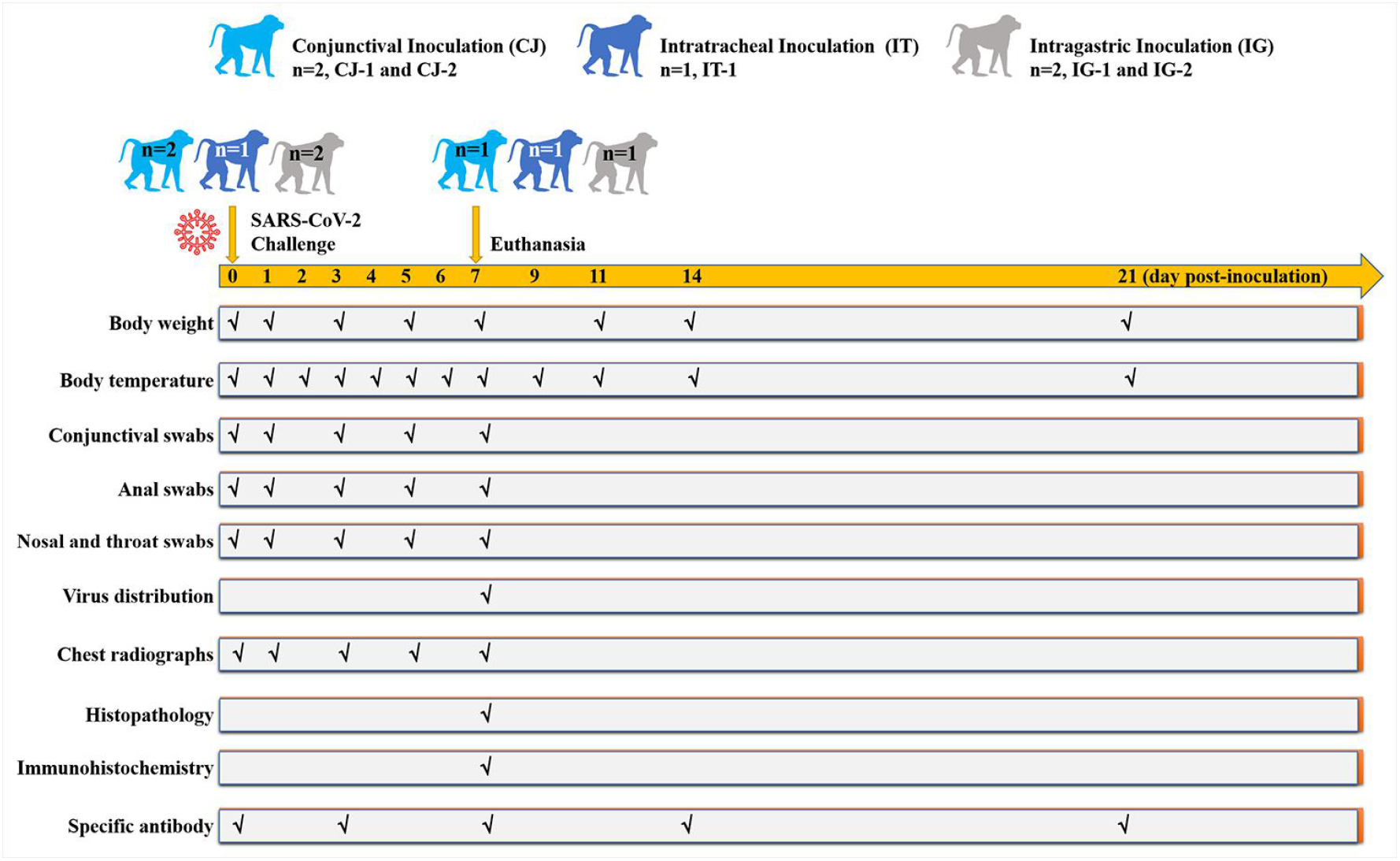
Graphic outline of experimental design and sample collection. Five male rhesus macaques (*Macaca Mulatta*) between the ages of 3 and 5 years were inoculated with 10^6^ TCID_50_/ml SARS-CoV-2. Two rhesus macaques via ocular conjunctival route named CJ-1 and CJ-2, one was inoculated via intratracheal route named IT-1, two were inoculated via intragastric route in sequence named IG-1 and IG-2, respectively. The macaques were observed daily for clinical signs (body weight temperature were tested as shown). On 0, 1, 3, 5, and 7 dpi, the conjunctival, nasal, throat and anal swabs were collected. CJ-1, IT-1, and IG-1 were euthanized and necropsied on 7 dpi. Tissues were collected to analysis the virus distributions. All sera were collected on 0, 7, 14 and 21 dpi for serologic detection to exam the SARS-CoV-2 specific IgG antibodies.

We daily observed the macaques for clinical signs. There was no significant change in the body weight (Figure 2A) in the CJ and IG inoculated macaques and the temperature (Figure 2B) in all the inoculated macaques, while the weight loss was obvious in the IT inoculated macaques and dropped about 125 g at 5 dpi. Routine specimens, including nasal and throat swabs, were collected on 0, 1, 3, 5, and 7-day post-inoculation (dpi). Additionally, to explore the potentially excretory routes of SARS-CoV-2 in the host, the conjunctival and anal swabs were also gathered.

**Fig. 2.**
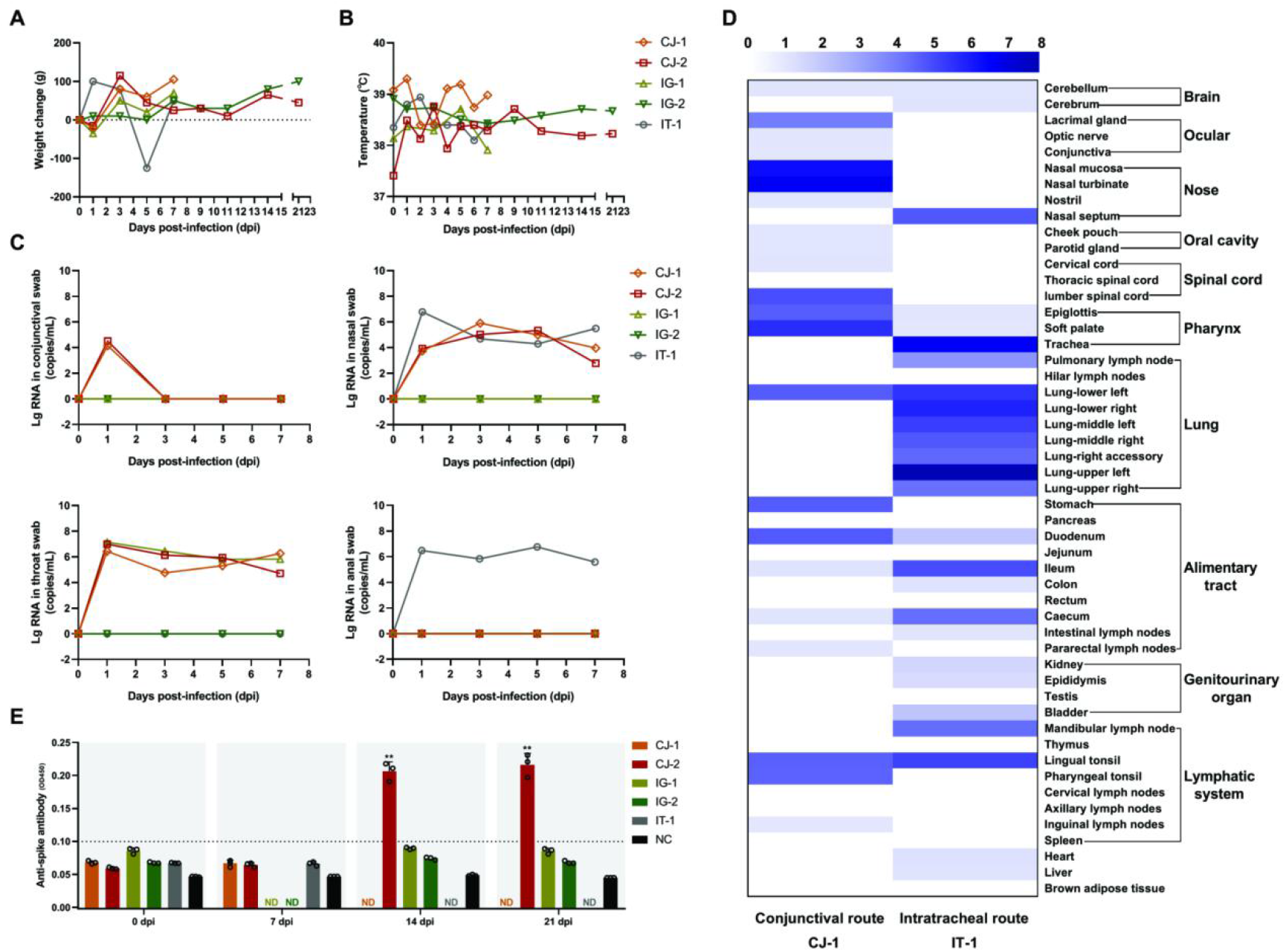
Clinical features, viral distributions and antibody detection in the sera from the rhesus macaques inoculated with SARS-CoV-2 via three routes. Clinical signs including body weight (A) and temperature (B) were observed. The viral load of the conjunctival, nasal, throat, and anal swabs specimens (C) from the five inoculated macaques on 0, 1, 3, 5, and 7 dpi. The comparison of viral distributionsin the majority of organs and tissues (D) from CJ-1 and IT-1 on 7 dpi. The darker the blue color, the higher the viral load. The specific IgG antibody against SARS-CoV-2 in the sera of the inoculated-macaques were tested by ELISA on 0, 7, 14, and 21 dpi (E). According to unpaired Welch’s *t*-test, the specific IgG antibody in the sera of conjunctival inoculated macaque exhibited a significant increase compared with prior to inoculation *(**p=0.0027)* and 21 dpi *(**p=0.0039)*. CJ-1 and CJ-2 were the two macaques that inoculated via conjunctival route, IT-1 was the macaque that inoculated via intratracheal route. IG-1 and IG-2 were the two macaques that inoculated via intragastric route. ND, not detected. NC, negative control (unpaired Welch’s *t*-test, \*\**p<0.01*).

Specifically, CJ and IT inoculated animals were able to detect a continued viral load in their nasal and throat swabs from 1 to 7 dpi. In contrast, the virus was not detected in any swabs from the IG inoculated macaques. Notably, only via the CJ route, viral load can be tested in conjunctival swabs (average, approximately 4.33 log_10_ RNA copies/mL) on 1 dpi and then became undetectable implying that the inoculated-SARS-CoV-2 may be transferred from the initial entry-conjunctiva to respiratory tract and other tissues. Meanwhile, only via the IT route, viral load can be ongoing examined in the anal swabs on 1-7 dpi and reached the peak about 6.76 log_10_ RNA copies/mL on 5 dpi, revealing distinct excretory pathways of host after inoculation via different routes (Figure 2C).

To determine the distribution of virus and histological lesions, CJ-1, IT-1, and IG-1 were euthanized and necropsied on 7 dpi. For CJ-1, viral load was primarily distributed in the nasolacrimal system and ocular, including the lacrimal gland (3.93 log_10_ RNA copies/mL), optic nerve (1.03 log_10_ RNA copies/mL), and conjunctiva (1.01 log_10_ RNA copies/mL); in the nose, including the nasal mucosa (6.25 log_10_ RNA copies/mL), nasal turbinate (6.63 log_10_ RNA copies/mL), and nostril (1.03 log_10_ RNA copies/mL); in the pharynx including epiglottis (4.57 log_10_ RNA copies/mL), soft palate (5.60 log_10_ RNA copies/mL); in the oral cavity including check pouch (1.03 log_10_ RNA copies/mL) and parotid gland (1.04 log_10_ RNA copies/mL); as well as in other tissues including lower left lobe of lung (4.6 log_10_ RNA copies/mL), tonsil (4.51 log_10_ RNA copies/mL), inguinal (1.01 log_10_ RNA copies/mL) and pararectal (6.25 log_10_ RNA copies/mL) lymph node, stomach (4.61 log_10_ RNA copies/mL), duodenum (4.66 log_10_ RNA copies/mL), ileum (1.08 log_10_ RNA copies/mL) and caecum (1.06 log_10_ RNA copies/mL) (Figure 1d). By contrast, for IT-1, the distribution of virus might be somewhat different owing to viral replication was highly in different lobes of the lung (10^4.22^ to 10^7.81^ copies/mL), and viral load was also widely detected in the nasal septum (4.69 log_10_ RNA copies/mL), tracheas (6.61 log_10_ RNA copies/mL), mandibular lymph node (4.26 log_10_ RNA copies/mL), tonsil (5.17 log_10_ RNA copies/mL), pulmonary lymph node (3.39 log_10_ RNA copies/mL), and some segments of the alimentary tract including duodenum (1.97 log_10_ RNA copies/mL), ileum (4.93 log_10_ RNA copies/mL), colon (1.11 log_10_ RNA copies/mL), and caecum (4.25 log_10_ RNA copies/mL) (Figure 2D). The distinct distributions of viruses by different inoculation routes were consistent with the anatomical structure, suggesting that the dispersion of SARS-CoV-2 into the host associated with the infection route. Furthermore, viruses were detectable in different segments of the alimentary canal and some lymphatic tissues via both inoculation routes revealing that they may play important roles in the spread of the virus within the host^5^. There were no significant histopathological changes in IG-1. Furthermore, the specific IgG antibody against SARS-CoV-2 was detectable in the CJ-2 at 14 and 21 dpi compared to before infection proofing that the animal was infected with SARS-CoV-2 (Figure 2E).

Meanwhile, comparison with prior to infection (day 0), the chest radiographs from CJ-1 revealed obscure lung markings and opaque glass sign in the bilateral upper lobes and the right lower lobe of the lung on 7 dpi. By comparison, IT-1 developed obviously increased radiographic changes on 7 dpi, exhibiting vessel convergence sign, obscure lung markings, marked ground-glass opacities, obscures costophrenic angle in the bilateral lobes of the lung, and patchy lesions in the right lower lobe of the lung (Figure 3A). Consistent with the radiographic alteration, microscopically, local lesions in the lungs from the CJ-1 displayed mild interstitial pneumonia characterized by widened alveolar interstitium, infiltration of inflammatory cells primarily including lymphocytes and monocytes, a small amount of exudation in the alveolar cavities (Figure 3B). Virus antigen was further confirmed by the SARS-CoV-2-specific antibody via immunohistochemistry (IHC) stain. In the damaged lobe of lungs, SARS-CoV-2 was predominantly observed in the alveolar epithelium and exfoliated-degenerative cellular debris in the alveolar cavities (Figure 3B) both in CJ-1 and IT-1. Notably, the results of IHC were highly in accordance with the data of viral load detection on 7 dpi both in the CJ-1 and IT-1. Specifically, viral antigens were scattered in several cells in the nasolacrimal system via conjunctival inoculation, while they were more prominent in the trachea via intratracheal inoculation (Figure 4). Moreover, viral antigen was obviously observed in the lamina propria of alimentary tract suggesting that the virus may spread from the initial entry to gut-associated lymphoid tissue (Supplementary Figure 1), and they were relatively slight detected in the kidney, myocardium, and liver in IT-1 but undetectable in CJ-1. (Supplementary Figure 2). Briefly, animal inoculated via conjunctival route gave evidence of relatively mild and local interstitial pneumonia compared to the intratracheal inoculated-macaque which demonstrated moderate, diffuse lesions in the lung accompanied with more infiltration of inflammatory cells and accumulation of exudation in the alveolar cavities.

**Fig. 3.**
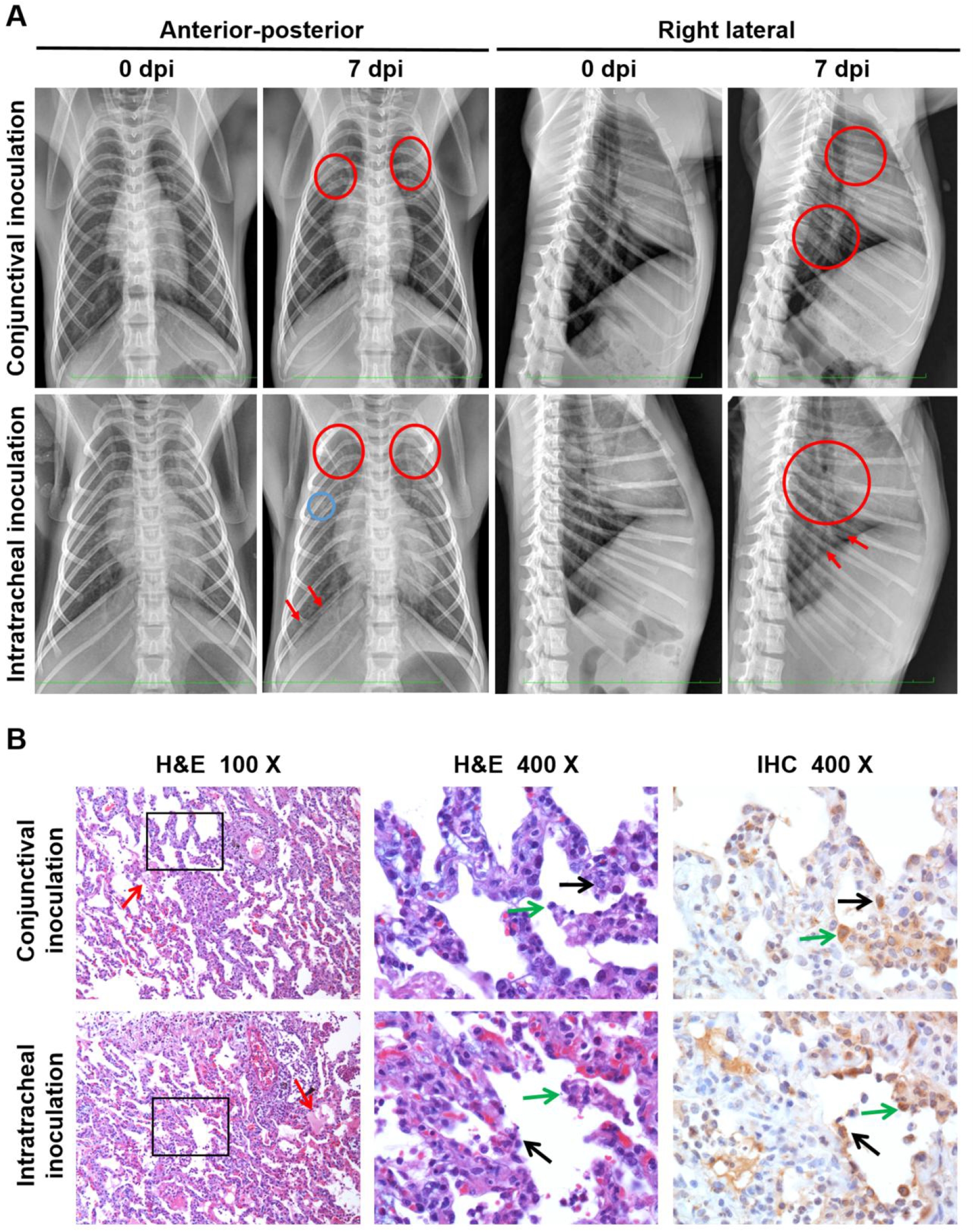
Compare the lesions in lungs from CJ-1 and IT-1 by radiographic alterations, histopathological and Immunohistochemical observation. The anterior-posterior and right lateral chest radiographs (A) from rhesus macaque imaged prior to SARS-CoV-2 inoculation (day 0) and 7 dpi. Areas of interstitial infiltration, indicative of pneumonia, are highlighted (red circle); obscures costophrenic angle (red arrows); patchy lesions (blue circle). Positional indicators are included (R=right). The histopathological and immunohistochemical observations in the lungs (B). Both the two macaques exhibited interstitial pneumonia with thickened alveolar septa, filtration of inflammatory cells mainly including lymphocytes and macrophages, some amounts of exudation (red arrows) in the alveolar cavities on 7dpi. Conjunctival route caused relatively mild pneumonia. The sequential sections were stained by HE and IHC, respectively. The viral antigens were observed primarily in the alveolar epithelia (black arrows) and the detached-degenerative cellular debris (green arrows). The H&E stained-sections under 400 magnification were the fractionated gain (black frame) of these sections under 100 magnification. The IHC section showed the same field with the black frame section under 400 magnification. Black scale bar = 100 μm, red scale bar = 50 μm.

**Fig. 4.**
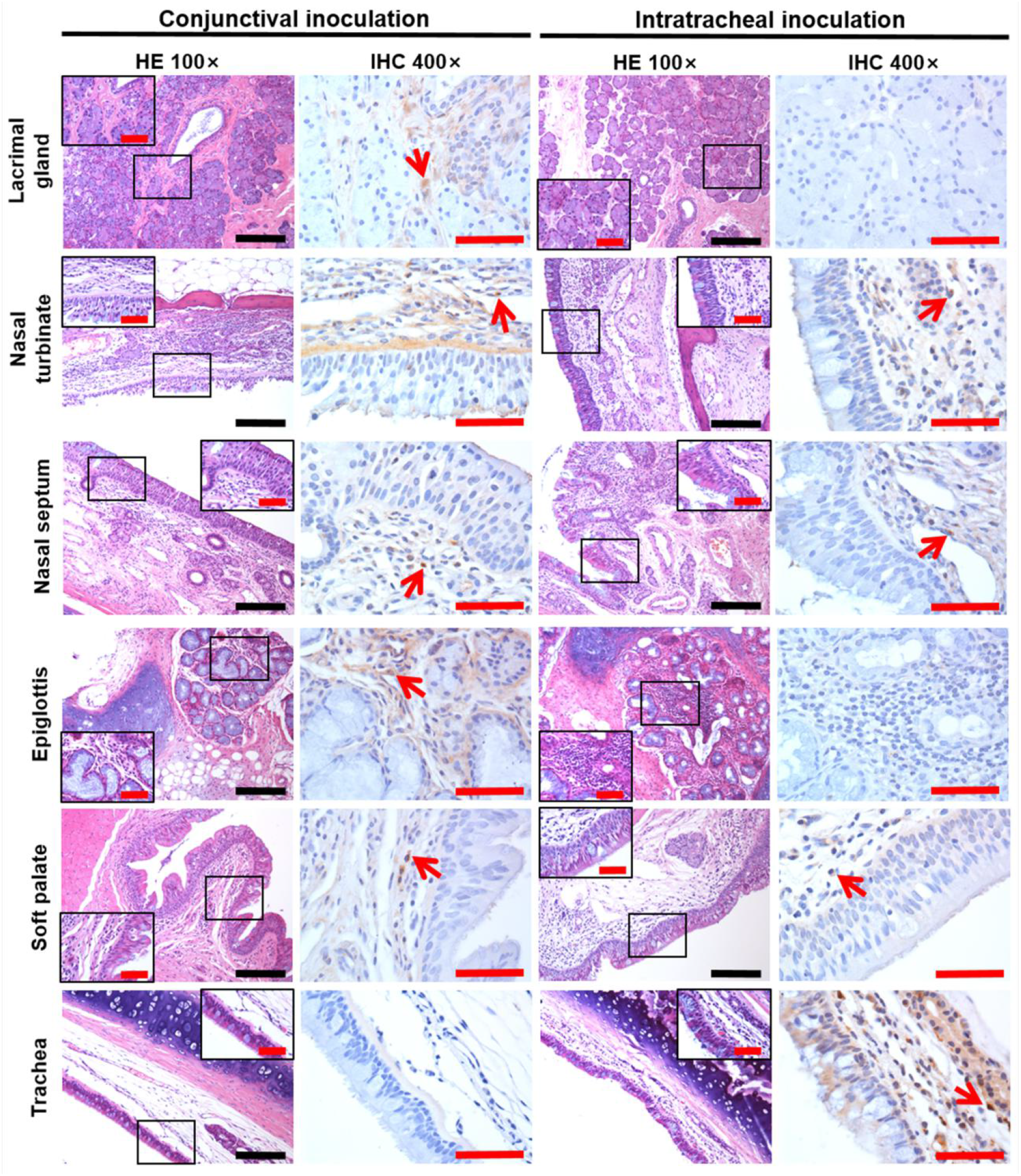
The distinct distributions of viruses by different inoculation routes between CJ-1 and IT-1 were consistent with the anatomical structure. The organs and tissues from the nasolacrimal system and the conductive system of the respiratory tract, including lacrimal gland, nasal turbinate, nasal septum, epiglottis, soft palate, and trachea were examined. The sequential sections were stained by HE and IHC, respectively. The field in the black frame was the fractionated gain of the HE slide. The IHC section showed the same field with the black frame section under 400 magnification. Black scale bar = 100 μm, red scale bar = 50 μm.

These data demonstrated that macaques can be infected with SARS-CoV-2 via the conjunctival route rather than the intragastric route. Compared to the intratracheal route, viral load was comparatively high in the nasolacrimal system and lesions in the lung were relatively mild and local via the conjunctival route. Similarly, both the two routes can affect the alimentary canal.

## Discussion

We inoculated rhesus monkeys via a single route via conjunctiva, intra-stomach, or intra-trachea, to avoid multiple routes of co-inoculation for confirming the exact pathway of inoculation. These results suggest that conjunctiva is a potential portal for viral transmission. In our results, viral load was detectable in several nasolacrimal system associated-tissues, especially in the conjunctiva, lacrimal gland, nasal cavity and throat, which drew the outline of the anatomical bridge between ocular and respiratory tissues. Particularly, the lacrimal duct functions as a conduit to collect and transport tear fluid from the ocular surface to the nasal-inferior meatus, being convenient for the drainage of virus from ocular to respiratory tract tissues. Actually, the previous report had demonstrated that although virus-containing fluid can be taken up through the conjunctiva, sclera, or cornea, the majority of liquid including tear and secretions is drained into the nasopharyngeal space or swallowed; the lacrimal duct epithelia are also possible to contribute to the absorption of tear fluid. Our results were highly consistent with the anatomical features that viruses enter the host via the conjunctival route. On the other hand, previous research demonstrated that SARS-CoV was undetectable in cynomolgus monkeys after intragastric inoculation which consistent with our result. At present, still, no evidence could prove that SARS-CoV-2 can transmission via fecal-oral route^5^, although viral RNA of SARS-CoV-2 was prolonged detectable in fecal samples from patients^6^ and in anal swabs from the infected macaques.

Respiratory viruses have the capability to stimulate ocular complications in infected patients, which then leads to respiratory infection ^7^. The fact that exposed mucous membranes and unprotected eyes increased the risk of SARS-CoV^1^ or SARS-CoV-2^2^ transmission suggests that healthcare professionals need to increase the awareness of eye protection in close contact with the patients or in crowded places.

## Methods

### Ethics statement

The animal biosafety level 3 (ABSL3) facility in the Institute of Laboratory Animal Science was used to complete all the experiments with rhesus macaques. All research was performed in compliance with the Animal Welfare Act and other regulations relating to animals and experiments. The Institutional Animal Care and Use Committee of the Institute of Laboratory Animal Science, Peking Union Medical College, reviewed and authorized all the programs in this research including animals (BLL20001).

### Cells and viruses

The SARS-CoV-2 named SARS-CoV-2/WH-09/human/2020/CHN was isolated by the Institute of Laboratory Animal Science, Peking Union Medical College. Vero cells were applied to the reproduction of SARS-CoV-2 stocks. Dulbecco’s modified Eagle’s medium (DMEM, Invitrogen, Carlsbad, USA) were applied to incubate this cell line at 37°C, 5% CO_2_, complemented with 10% fetal bovine serum (FBS), 100μg/ml streptomycin, and 100 IU/ml penicillin, and maintained. Titers for SARS-CoV-2 were resolved by TCID_50_ assay.

### RNA extraction and RT-PCR

All the collected-organs were applied to extract Total RNA as the description in the previous report. Briefly, the RNeasy Mini Kit from Qiagen, Hilden, Germany and the PrimerScript RT Reagent Kit from TaKaRa, Japan were used following manufacturer instructions. RT-PCR reactions were applied to the PowerUp SYBG Green Master Mix Kit from Applied Biosystems, USA, following cycling protocol: 50°C for 30 min, followed by 40 cycles at 95°C for 15 min, 94°C for 15 s, and 60°C for 45 s. The primer sequences used for RT-PCR were targeted against the envelope (E) gene of SARS-CoV-2. The forward primer is 5’-TCGTTTCGGAAGAGACAGGT-3’, the reverse primer is 5’-GCGCAGTAAGGATGGCTAGT-3’.

### Animal experiments

Five male rhesus macaques (*Macaca Mulatta*) between the ages of 3 and 5 years were used in this research. All were negative for tuberculosis and simian immunodeficiency virus. They were inoculated with 1×10^6^ 50% tissue-culture infectious doses (TCID_50_) of SARS-CoV-2 via three routes. Two male rhesus macaques were inoculated with 10^6^ TCID_50_/ml SARS-CoV-2 via ocular conjunctival route named CJ-1 and CJ-2, one was inoculated via intratracheal route named IT-1, two were inoculated via intragastric route in sequence named IG-1 and IG-2, respectively. Prior to sample collection, all animals were anesthetized with ketamine hydrochloride (10 mg/kg). On 1, 3, 5, and 7 dpi, the conjunctival, nasal, throat and anal swabs were collected and incubated in 1 ml DMEM with 50 μg/ml streptomycin and 50 U/ml penicillin. The IPTT-300 temperature probes, which were injected interscapular into the macaques prior to the start of the experiment, were applied to do temperature monitor every day. CJ-1, IT-1, and IG-1 were euthanized and necropsied on 7 dpi. Tissues were collected as followed, conjunctiva, lacrimal gland, optic nerve, cerebellum, cerebrum, different segments of the spinal cord, nostril, nasal turbinate, nasal mucosa, nasal septum, soft palate, cheek pouch, parotid gland, epiglottis, lingual tonsil, pharyngeal tonsil, different lobes of lung, trachea, different lymph nodes, heart, liver, spleen, pancreas, different segments of the alimentary canal, kidney, bladder, testis, and brown adipose tissues samples for detecting the viral loads to analysis the virus distributions. All sera were collected on 0, 7, 14 and 21 dpi for serologic detection to exam whether there presence the IgG antibodies reactive with SARS-CoV-2 antigens.

### Preparation of Homogenate Supernatant

An electric homogenizer was applied to prepare tissues homogenates by 2 min 30s incubated in 1ml of DMEM. The homogenates were centrifuged at 3,000 rpm at 4°C for 10 min. The supernatant was harvested and reposited for viral titer at −80°C.

### ELISA antibody assay

Sera of Each animal were collected to detect the SARS-CoV-2 antibody through enzyme-linked immunosorbent assay (ELISA) on 0, 7, 14 and 21 dpi. The 96-well plates coated with 0.1μg Spike protein of SARS-CoV-2 from Sino Biological (Product code: 40591-V08H) at 4°C overnight were blocked by 2% BSA/PBST at room temperature for 1 hour. Sera samples were diluted at 1:100, and then were added to different wells and maintained at 37 °C for 30 minutes, followed by the goat anti-monkey antibody labeled with horseradish peroxidase (Abcam, ab112767) incubated at room temperature for 30 minutes. The reaction was determined at 450 nm.

### Hematoxylin and Eosin Staining

Ten percent buffered formalin solution fixed all the collected organs, and paraffin sections (3-4μm in thickness) were prepared according to routine practice. All the tissue sections were stained with Hematoxylin and Eosin. The histopathological changes of different tissues were observed under an Olympus microscope.

### Immunohistochemistry (IHC)

Ten percent buffered formalin solution fixed all the collected organs, and paraffin sections (3-4 μm in thickness) were prepared routinely as the description in the previous report. Briefly, an antigen retrieval kit (AR0022, Boster) was applied to the sections at 37°C for 1 min. Three percent H_2_O_2_ in methanoland were fulfilled to quench for endogenous peroxidases for 10 min. The slices were incubated at 4°C overnight with a laboratory prepared-7D2 monoclonal antibody after blocking in one percent normal goat serum. HRP-labeled goat anti-mouse IgG secondary antibody from ZDR-5307, Beijing ZSGB Biotechnology were maintained at 37°C for 60 min. The 3,30-diaminobenzidine tetrahydrochloride was treated to make the results visual. The sections were counterstained with hematoxylin, dehydrated and mounted on a slide and observed by the Olympus microscope. The sequential sections from all collected tissues were directly incubated with HRP-labeled goat anti-mouse IgG were used as the omission control of viral antigen staining. The sequential sections from all collected tissues were incubated with a recombinant anti-Mouse IgG antibody [RM104] (ab190475, Abcam) performed as the negative control for the expression of viral antigen.

### Statistical analysis

All data were analyzed by GraphPad Prism 8.0 software. Statistically significant differences were determined using Welch’s *t*-tests. A two-sided p value < *0.05* was considered statistically significant. \**P* < 0.05, \*\**P* < 0.01, \*\*\**P* < 0.001.

## Acknowledgments

This work was supported by the National Key Research and Development Project of China (Grant No. 2016YFD0500304), CAMS initiative for Innovative Medicine of China (Grant No. 2016-I2M-2-006), National Mega projects of China for Major Infectious Diseases (Grant No. 2017ZX10304402).

## Authors Contributions

Conceptualization: C.Q.; Methodology:C.Q., W.D., L.L.B., H.G., Z.G.X., Y.J.Q., Z.Q.S.,; Investigation: W.D., L.L.B., H.G., Z.G.X., Y.J.Q., Z.Q.S., S.R.G., J.Y.L., J.N.L., P.Y., F.F.Q., Y.F.X., F.L.L., C.X., Q.L., J.X., Q.W., M.Y.L., G.P.W., S.Y.W., H.S.Y., X.L., W.J.Z., Y.L.H., C.Q.; Writing – Original Draft: Z.Q.S. and J.X.; Writing–Review and Editing: C.Q.; Funding Acquisition: C.Q. and L.L.B.,; Resources: C.Q.; Supervision: C.Q.

---

**Supplementary Figure 1.**
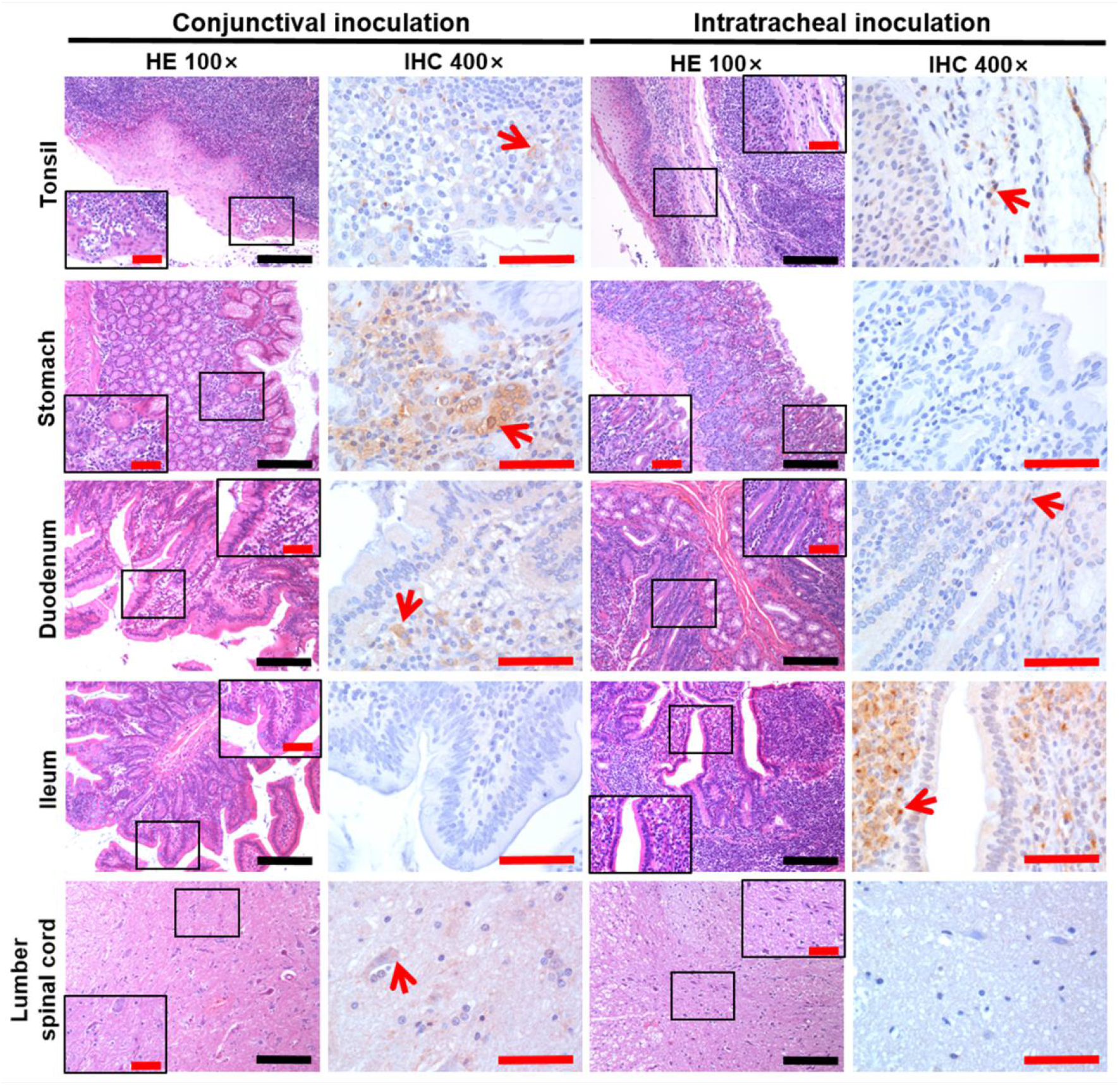
The comparison of viral distributions in the tonsil, the alimentary tract including stomach, duodenum, and ileum, and the lumber spinal cord between CJ-1 and IT-1. The sequential sections were stained by HE and IHC, respectively. The field in the black frame was the fractionated gain of the HE slide. The IHC section showed the same field with the black frame section under 400 magnification. Black scale bar = 100 μm, red scale bar = 50 μm.

**Supplementary Figure 2.**
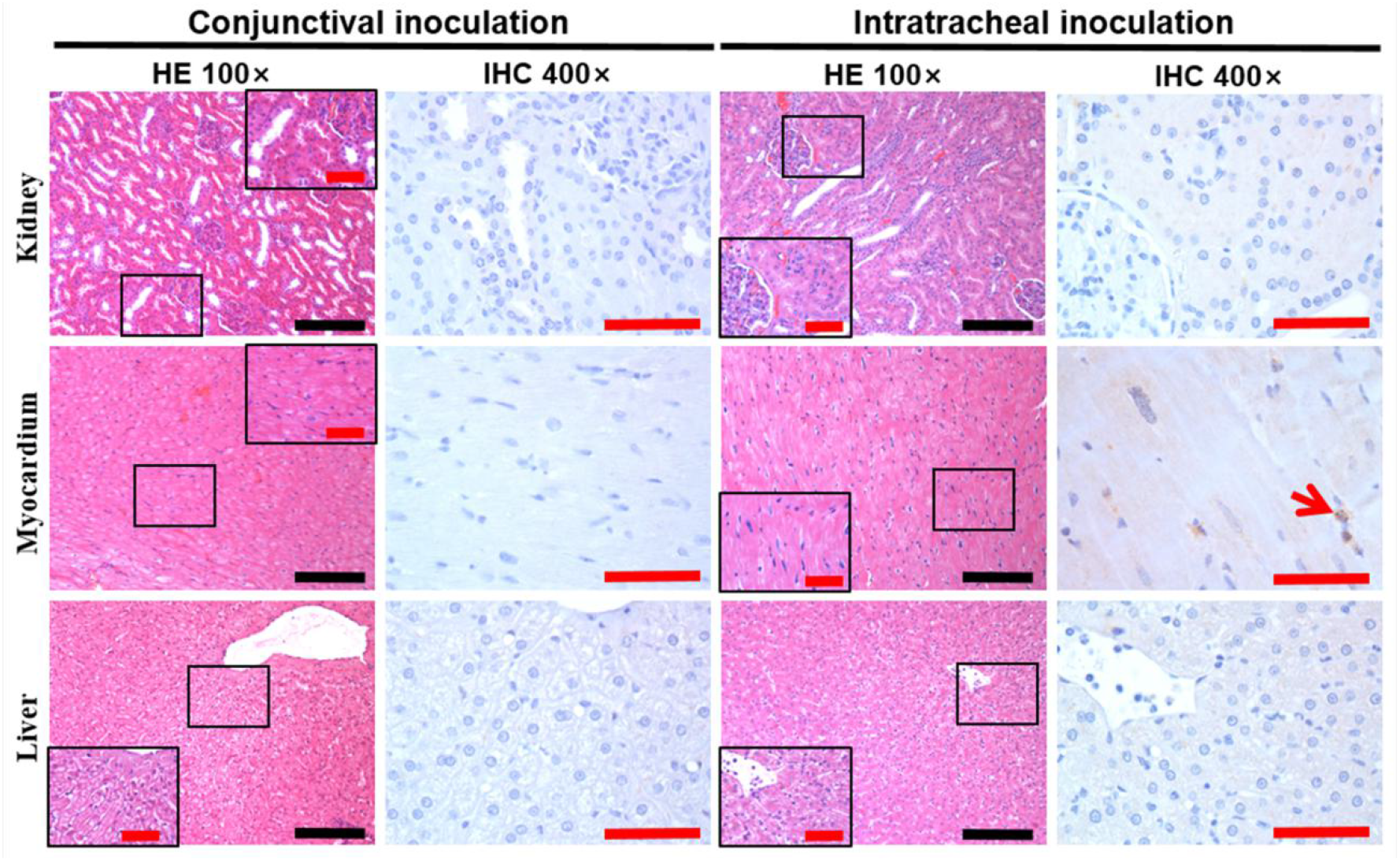
Slight viral antigen was detected in the kidney, heart, and liver in the IT-1 and that was negative in CJ-1. The sequential sections were stained by HE and IHC, respectively. The field in the black frame was the fractionated gain of the HE slide. The IHC section showed the same field with the black frame section under 400 magnification. The field in the red frame was the fractionated gain of the IHC slide. Black scale bar = 100 μm, red scale bar = 50 μm.

